# Spatial-temporal targeted and non-targeted surveys to assess microbiological composition of drinking water in Puerto Rico following Hurricane Maria

**DOI:** 10.1101/2021.05.07.442998

**Authors:** Maria Sevillano, Solize Vosloo, Irmarie Cotto, Zihan Dai, Tao Jiang, Jose M. Santiago Santana, Ingrid Y. Padilla, Zaira Rosario-Pabon, Carmen Velez Vega, José F. Cordero, Akram Alshawabkeh, April Gu, Ameet J. Pinto

**Affiliations:** Department of Civil and Environmental Engineering, Northeastern University, Boston, MA, USA; Key Laboratory of Drinking Water Science and Technology, Research Center for Eco-Environmental Sciences, Chinese Academy of Sciences, Beijing, China; University of Chinese Academy of Sciences, Beijing, China; Department of Natural Sciences, University of Puerto Rico, Carolina, PR, USA; Department of Civil Engineering and Surveying, University of Puerto Rico, Mayagüez, PR, USA; University of Puerto Rico—Medical Sciences Campus, San Juan, PR, USA; Department of Epidemiology and Biostatistics, University of Georgia, Athens, Georgia, USA; School of Civil and Environmental Engineering, Cornell University, Ithaca, NY, USA

**Keywords:** drinking water quality, hurricane Maria, metagenomics, qPCR, genome resolved metagenomics

## Abstract

Loss of basic utilities, such as drinking water and electricity distribution, were sustained for months in the aftermath of Hurricane Maria’s (HM) landfall in Puerto Rico (PR) in September 2017. The goal of this study was to assess if there was deterioration in biological quality of drinking water due to these disruptions. This study characterized the microbial composition of drinking water following HM across nine drinking water systems (DWSs) in PR and utilized an extended temporal sampling campaign to determine if changes in the drinking water microbiome were indicative of HM associated disturbance followed by recovery. In addition to monitoring water chemistry, the samples were subjected to culture independent targeted and non-targeted microbial analysis including quantitative PCR (qPCR) and genome-resolved metagenomics. The qPCR results showed that residual disinfectant was the major driver of bacterial concentrations in tap water with marked decrease in concentrations from early to late sampling timepoints. While *Mycobacterium avium* and *Pseudomonas aeruginosa* were not detected in any sampling locations and timepoints, genetic material from *Leptospira* and *Legionella pneumophila* were transiently detected in a few sampling locations. The majority of metagenome assembled genomes (MAGs) recovered from these samples were not associated with pathogens and were consistent with bacterial community members routinely detected in DWSs. Further, whole metagenome-level comparisons between drinking water samples collected in this study with samples from other full-scale DWS indicated no significant deviation from expected community membership of the drinking water microbiome. Overall, our results suggest that disruptions due to HM did not result in significant and sustained deterioration of biological quality of drinking water at our study sites.

**Graphical Abstract:** 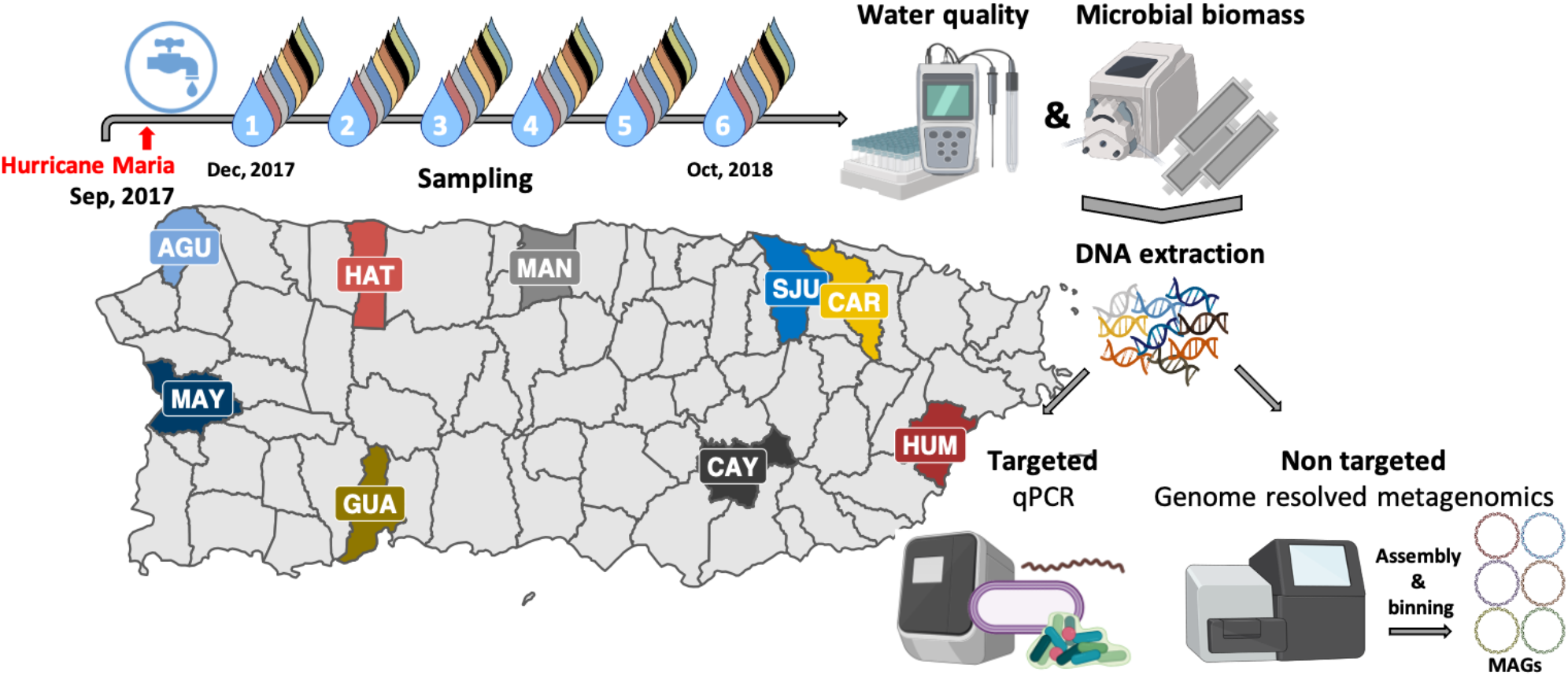

## Introduction

A 2015 report on the Safe Drinking Water Act violations in Puerto Rico (PR) indicated high levels of contaminants such as volatile organic compounds (VOC), total coliform bacteria, and disinfection by products (DBPs) impacted around 70% of the islands population(NRDC, 2017). This report recommended investment in drinking water systems (DWSs), including treatment, distribution system upgrade and maintenance, and source water protection. Such investments are also important across the US, as the water infrastructure continues to age(ASCE, 2017) and water quality violations are being increasingly reported(Allaire et al., 2018). Further complicating the issue of providing regulation compliant water while relying on an aging water infrastructure is the increasing frequency and intensity of extreme weather events(Estrada et al., 2015; Goodess, 2012). In the year 2017 alone, Hurricanes Harvey, Irma, and Maria (HM) caused widespread damages and were categorized as historic billion dollar disasters in the US(NOAA NCEI, 2020). The resiliency of DWSs during these extreme events is particularly important, as lack of access to safe drinking water may result in further detrimental health impacts. Natural disasters can contaminate source waters, impacting proper treatment, distribution, and ultimately affect consumer health(Ashbolt, 2015; Exum et al., 2018) Previous studies have highlighted water quality degradation associated with extreme weather events like hurricanes. Schwab et al. measured concentrations of faecal coliforms, E. coli, and enterococci in tap and surface waters following Hurricane Katrina and did not recover any of the bacterial indicators in tap water samples irrespective of chlorine residual concentration(Schwab et al., 2007). A recent study on the impacts of Hurricane Harvey on water quality from two DWSs in Texas highlighted that source water quality and water demand and their relationship to water age strongly impacted microbial communities and influenced the time for recovery(Landsman et al., 2019). Similarly, a amplicon-sequencing based study carried out in St. Thomas, post Hurricane Irma and HM revealed that the microbial community structure in rain cisterns, coastal stations, and surface runoff waters was dramatically different between sampling sites with faecal indicator bacteria (FIB) detected in cisterns used as household water supply(Jiang et al., 2020).

HM, classified as a category 4 hurricane, impacted 3 million people in PR. Loss of basic utilities (i.e., water, cellular coverage, and electricity) was associated with remoteness category(Kishore et al., 2018). Water services generally recovered quickly in densely populated areas, while remote areas either recovered quickly or months later. However, electricity services took longer to recover irrespective of remoteness category. Even when water services were restored, intermittent water supply was common due to unreliable electrical supply; this could potentially degrade water quality via stagnation and loss of disinfectant residual, intrusion, and backflows(Bautista-de los Santos et al., 2019). Boil advisories and point of use chlorination were in place after water services resumed and were reported by the Puerto Rico Aqueducts and Sewers Authority (PRASA) through mid-January 2018(Exum et al., 2018). Previously, Lin et al.(Lin et al., 2020) provided insights into metals, micropollutants, and molecular toxicity of pre- and post-HM drinking water samples in PR and showed the impact of HM on chemical water quality, suggesting that trace metals were potential drivers of cumulative risk from drinking water. Additionally, Keenum et al.(Keenum et al., 2021) characterized five unregulated small scale DWS and one large PRASA DWS in PR six months after HM. In this study, targeted culture and molecular based analyses (i.e., quantitative PCR (qPCR), 16S rRNA amplicon sequencing) demonstrated similar microbial communities and concentrations of opportunistic premises plumbing pathogens (OPPPs) compared to those reported in the continental US. In our study, we also aim to evaluate the microbial water quality in the aftermath of HM. However, unlike Keenum et al(Keenum et al., 2021), we conducted a recurrent sampling campaign beginning in December 2017 spanning nine locations across PR for a duration of a year. Despite the magnitude of HM in PR, there hasn’t been a large effort to characterize microbial water quality. To date, there have been two reports focused on chemical contamination(Lin et al., 2020; Warren, 2019) and two (including this one) on microbial composition(Keenum et al., 2021) of DWDs on the island. These studies are essential to establish relationships, sampling infrastructure, and methodologies needed to respond to future storms, as well as to communicate risk and execute corrective actions to decrease exposure risk and unwanted health outcomes. Thus, our goals were (1) to utilize an extended spatial-temporal sampling campaign to determine if changes in drinking water microbiome were indicative of disturbance followed by recovery, m(2) if this disturbance-recovery dynamic was associated with presence of potential pathogens, (3) whether potential pathogen presence was persistent or transient, and finally (4) whether microbial composition of PR drinking water was consistent with or deviated significantly from other drinking water systems.

## 2. Materials and methods

### 2.1 Drinking water sampling and water quality analyses

Nine sampling locations were chosen across different geographic locations in PR. Tap water was filtered in triplicate on site through 0.2 µm Sterivex filters (EMD Millipore™, Cat. no. SVGP01050) using a field peristaltic pump (Geotech, Cat. no. 91352123) until the filter clogged or up to a 20L volume for each filter. Water quality parameters (i.e., temperature, pH, conductivity, and dissolved oxygen) were recorded on site with an Orion Star probe (Thermo Scientific, Cat. no. 13645571). A portable spectrophotometer (HACH, Cat. no. DR1900-01H) was used to measure Total chlorine (HACH, Cat. no. 2105669) and phosphate (HACH, Cat. no. 2106069) on site. Nitrogen species (i.e., ammonia, nitrate, nitrite) were measured in the laboratory with a HACH spectrophotometer using HACH test and tube format (HACH, Cat. no. 2606945, 2605345, 2608345, respectively). Total Organic Carbon (TOC) was measured with a Shimadzu TOC-LCPH Analyzer (Shimadzu, Kyoto, Japan). Additional details about the 54 samples can be found in Table S1.

### 2.2 DNA extraction, qPCR and shotgun sequencing

DNA extractions were performed using a modified version of the DNeasy PowerWater Kit (QIAGEN, Cat no. 14900-50-NF) protocol(Vosloo et al., 2019). Briefly, the polyethersulfone (PES) membrane from the Sterivex filter was processed by aseptically cutting it into smaller pieces and transferring to a Lysing Matrix E tube (MP Biomedical, Cat. no. MP116914100). Subsequently, 294 µL of 10X Tris-EDTA buffer pH 8 (G-Biosciences, Cat. no. 501035446) was added to the Lysing Matrix E tube and supplemented with 6 µL of lysozyme (50 mg mL^-1^, Thermo Fisher Scientific, Cat. no. 90082), followed by a 60 min incubation at 37°C with mixing at 300 rpm. Subsequently, 300 µL of PW1 solution from DNeasy PowerWater Kit was mixed in and 30 µL of Proteinase K (20 mg mL^-1^, Fisher Scientific, Cat. no. AM2546) was added. An incubation period of 30 min at 56°C with mixing at 300 rpm followed. Previously removed spheres from the corresponding Lysing E matrix tube were replenished and 630 µl chloroform: isoamyl alcohol (Fisher Scientific, Cat. no. AC327155000) was added. Bead beating was performed at setting 6 for 40 seconds using a FastPrep – 24™ (MP Biomedical, Cat. no. 116004500). The resulting homogenized mixture was centrifuged for 10 min at 14,000 x g at 4°C and the upper aqueous phase was transferred to a clean 1.5 mL tube. A supplement of 6 µL carrier RNA (prepared by mixing 310 µl of Buffer EB from DNeasy PowerWater Kit with 310 µg lyophilized carrier RNA (QIAGEN, Cat. no. 1068337)) was mixed with 600 µL of recovered supernatant. This was then purified using the automated DNA purification protocol with DNeasy PowerWater Kit on a QIACube system (QIAGEN, Cat. no. 9001292). In addition to the samples, controls were processed identically and consisted of unused transported Sterivex filter membranes (filter blank), no input material (reagent blank), and sterilized deionized water filtered through Sterivex (water blank). This set of three controls were included with each sampling campaign (n=6) and extraction run.

qPCR was performed on a QuantStudio™ 3 Real-Time PCR System (ThermoFisher Scientific Cat. no. A28567). PCR reactions were carried out in a 20 µl volume containing Luna Universal qPCR Master Mix (New England Biolabs, Inc., Cat. no. NC1276266), F515 (GTGCCAGCMGCCGCGGTAA) and R806 (GGACTACHVGGGTWTCTAAT) primer pair(Caporaso et al., 2011) (IDT), DNAse/RNAse-Free water (Fisher Scientific, Cat. no. 10977015), and 5 µL of 10X diluted DNA template. Reactions were prepared by an epMotion M5073 liquid handling system (Eppendorf, Cat. no. 5073000205D) in triplicate. The cycling conditions were as follows, initial denaturing at 95 °C for 1 min followed by 40 cycles of denaturing at 95 °C for 15 s, annealing at 50 °C for 15 s, and extension 72 °C for 1 min. Melting curve analyses was performed by ramping from 72 °C to 95 °C for 15 s, and 60 °C for 1 min, 95 °C for 15 s. A negative control (NTC) and a standard curve consisting of 7 points ranging from 10^1^-10^7^ copies of 16S rRNA gene were included in every qPCR run. Genomic DNA from selected samples were sent to University of Illinois Roy J. Carver Biotechnology Center (UI-RJCBC) for library preparation using a low input DNA kit (NuGEN, Cat. no 0344NB). Libraries were loaded into two SP lanes on a NovaSeq 6000 instrument with an output of 2×150nt reads. The prepared libraries included 33 samples and 3 pooled blanks (i.e., filter blank, reagent blank, and water blank) from all locations.

### 2.3 qPCR analyses for waterborne pathogens

Previously published primers targeting *Legionella* spp.(Nazarian et al., 2008; Yáñez et al., 2005), *Mycobacteria* spp.(Chern et al., 2015; Radomski et al., 2010), pathogenic Leptospira(Stoddard et al., 2009), and *Pseudomonas aerugionosa*(Anuj et al., 2009) were used for qPCR assays. Reactions were set up by an epMotion M5073 liquid handling system in triplicate. The assays consisted of 2X PrimeTime Gene Expression Master Mix (IDTDNA, Cat no. 290479057) with low reference ROX dye, target primers and probe (IDTDNA), 10-fold diluted template and water (UltraPure™ DNase/RNase-Free Distilled Water, Thermo Fisher Scientific, Cat. no. 10977015). Single target reactions were conducted in a total volume of 20 µL, whereas duplex qPCR (i.e., *Pseudomonas aeruginosa* assay) were conducted in 25 µL. Primer and probe sequences and cycling conditions are described in Table S2. Standard cycling conditions for all reactions were programmed on a QuantStudio™ 3 Real-Time PCR System. Target gene copy numbers were determined by comparing threshold cycle with standard curve generated using gblocks gene fragments as standards (Table S2).

### 2.4 Metagenomic reads pre-processing

The raw reads obtained from UI-RJCBC were processed with fastp(Chen et al., 2018) v0.19.7 to remove homopolymer stretches using the following flags ‘--trim_poly_g --trim_poly_x’. Further, trimmed reads were mapped with BWA-MEM(Li, 2013) v0.7.12 against the UniVec database build 10.0 (National Center for Biotechnology Information 2016) to perform vector screening and retained unmapped paired reads. Subsequently, Nonpareil(Rodriguez-R et al., 2018) v3.303 was used on the quality filtered reads to estimate average community coverage and metagenomic dataset diversity using kmer algorithm with a kmer size of 20 and default parameters.

### 2.5 Metagenome assembly and mapping

Sample reads were co-assembled based on each sampling location using metaSPAdes(Nurk et al., 2017) v 3.11.1 with the following flags ‘-meta -t 16 --phred-offset 33 -m 500 -k 21,33,55,77,99,119’ and further filtered to a minimum scaffold length of 500 bp. Reads from samples and controls (i.e., extraction blank, filter blank, DI water blank) were mapped to co-assemblies using BWA-MEM. An approach similar to Dai et al.(Dai et al., 2020) was used to remove potential contaminant scaffolds. Briefly, BWA-MEM with flag -F4 and -f2 was used to map sample and control reads against co-assemblies. The BEDtools(Quinlan and Hall, 2010) genomecov using flags -g, and -d was used to calculate coverage and per base coverage using generated BAM files. Relative abundances (RA) and normalized coverage deviation (NCD) were calculated for each scaffold with coverage and per base coverage information, respectively. Scaffolds that were not detected in the blanks or for which sample RA was greater than the blank RA and the sample NCD was less than the blank NCD were considered true scaffolds and were used in downstream analyses. All scaffolds that did not meet these criteria were considered contaminant scaffolds and removed from further analyses. Assembly statistics were obtained from QUAST(Gurevich et al., 2013) 5.0.2. To contextualize the metagenome assemblies with respect to other distribution systems, assemblies from other drinking water systems (considered unperturbed systems because samples were not associated with any natural disaster or water quality issues) were compared against our co-assemblies. MASH(Ondov et al., 2016) v2.1.1 was used to estimate the dissimilarity between assemblies using ‘-r’ and ‘-m 2’ flags, and a sketch size of 100000.

### 2.6 Taxonomic classification of metagenomic assemblies

The taxonomic classification of scaffolds was performed with a contig annotation tool(Von Meijenfeldt et al., 2019) (CAT v5.0.4) program in which open reading frames (ORF) are predicted with Prodigal(Hyatt et al., 2010) and used as alignment queries by DIAMOND(Buchfink et al., 2014) against the NCBI non-redundant (nr) protein database (downloaded ftp://ftp.ncbi.nlm.nih.gov/blast/db/FASTA/, 2020-03-04). Selected genera known to contain pathogenic species, as well as non-pathogenic species, that are relevant to drinking water systems were further examined. The CAT annotations of true scaffolds were evaluated against annotated controls and only scaffolds with rpoB normalized coverage above controls and additional annotation support were considered to avoid false positives at the genus level. Specifically, additional support consisted of classification with kaiju(Menzel et al., 2016) v1.7.2 using reference indexes containing NCBI BLAST nr database and microbial eukaryotes with default parameters (downloaded http://kaiju.binf.ku.dk/server,nr_euk 2019-06-25) and/or by kraken2(Wood et al., 2019) v2.0.9-beta against RefSeq database (downloaded https://lomanlab.github.io/mockcommunity/mc_databases.html). Additionally, we examined the scaffolds of eukaryotic origin with metaEuk(Levy Karin et al., 2020) v3.8dc7e0b to assign taxonomy using a publicly available MMseqs2 database containing protein profiles from the marine eukaryotic reference catalogue (MERC), Marine Microbial Eukaryote Transcriptome Sequencing Project (MMETSP), and Uniclust50 (downloaded http://wwwuser.gwdg.de/~compbiol/metaeuk/2020_TAX_DB/).

### 2.7 Metagenomic assembled genomes

Co-assemblies were binned with CONCOCT(Alneberg et al., 2014) within anvi’o(Eren et al., 2015) v5.1 by clustering scaffolds 2500 bp or longer into metagenome assembled genomes (MAGs) and manually refining them within the anvi’o platform. Furthermore, dRep(Olm et al., 2017) v2.3.2 was used to dereplicate MAGs and obtain representative genomes with flags ‘-comp 50 -con 10’ and default values. GTDB-tk(Parks et al., 2018) v0.1.3 was used to assign taxonomy to MAGs with the flag ‘classify_wf’. Sample reads were mapped to corresponding single pseudo contig MAGs. Pseudo contigs were generated with the union command in EMBOSS(Rice et al., 2000) utility. Mapping was performed with BBMap(Bushnell, 2015) v38.24 using a 90% identity threshold and setting flags ‘ambiguous=best’, ‘mappedonly=t’, and ‘pairedonly=t’. Detection of a MAG in a sample was established when ≥ 25% of its bases were covered by at least one read from the corresponding sample. Coverage was determined with samtools(Li et al., 2009) v1.10 ‘coverage’. The abundance of a MAG in a sample was calculated as sample reads mapped per million reads per genome length in kbp (RPKM). Further information about MAGs, such as number of 5S rRNA, 16S rRNA, 23S rRNA, and tRNA counts was obtained by annotating the MAGs using DRAM(Shaffer et al., 2020) v1.0.6. The databases used with DRAM were downloaded with the following flags ‘DRAM-setup.py prepare_databases --output_dir DRAM_data --skip_uniref’. MAGs from this study were compared to 52,515 MAGs recovered from environmentally diverse metagenomes by Nayfach et al.(Nayfach et al., 2020) (downloaded https://portal.nersc.gov/GEM/genomes/fna.tar, 2020-11-10) using FastANI(Jain et al., 2018) v2.3.2 with default parameters. Metadata linked with this genomic catalogue of earth microbiomes, hereafter referred to as JGI MAGs (downloaded https://portal.nersc.gov/GEM/genomes/genome_metadata.tsv, 2020-11-30) was used to address niche association. Further, SEARCH-SRA(Stewart et al., 2015; Torres et al., 2017; Towns et al., 2014) online portal was used to interrogate the SRA database (246,329 records) by aligning metagenomic datasets to our MAGs. Only records that mapped 10 or more reads from the SRA collection were further inspected. The metadata associated with SRA accession numbers (downloaded https://s3.amazonaws.com/starbuck1/sradb/SRAmetadb.sqlite.gz, 2021-04-08) was incorporated through custom scripts in R software(R Development Core Team, 2016) that rely on the dbplyr(Wickham et al., 2021) package. Records that were classified as retrieved from metagenomic library sources and with a whole genome sequence strategy were retained for analysis. Considering that SRA metadata is user provided, manual curation to ensure consistency and retrieve the same ecosystem categories as in JGI MAGs metadata was performed. Data with missing context (lacking information in title or description) were considered as “others” and removed from analyses. The association between a MAG and an ecosystem category was determined by multiplying the total number of reads from the ecosystem category mapping to the MAG by the ratio of the number of unique SRA records associated with a MAG and ecosystem category and the number of unique SRA records within the entire dataset assigned to the ecosystem category.

### 2.8 Data analyses and statistics

Statistical analyses were conducted in R and visualizations generated with ggplot2(Wickham, 2011) package. PCA analyses of water quality parameters were performed with centered and scaled data in R base prcomp(). Linear regression models of log10 (volume normalized 16S rRNA gene copies) against water quality parameters were fit using base R lm(). Pearson correlation between log10(volume normalized 16S rRNA gene copies) and chlorine concentrations (mg/L) was obtained with base R cor() and exponential decay curve with nls(). Euclidian distances between pairwise Mash distance of DW metagenomes was calculated with vegan(Oksanen et al., 2015) function vegdist() and clustered with complete linkage method with base R hclust(). PCoA ordination of Mash distances was performed with ape(Paradis et al., 2004) function pcoa(). Group wise non parametric testing was performed with R base statistic packages using function kruskal.test() or wilcox.test(). Permutational hypothesis testing (n iterations=10000) of differences in group means between two groups was performed after up-sampling group data to balance observations using upsample() from the groupdata2(Olsen, 2021) package. Analyses of variance (ANOVA) was performed with aov() and followed up with post hoc Tukey-Kramer testing using TukeyHSD() in base R.

## 3. Results and discussion

### 3.1 Bacterial concentrations were associated with water quality parameters, particularly total chlorine concentrations

Water quality parameters were recorded for each sampling location and timepoint (Figure1A, Table S1). PCA analyses was conducted to assess whether water chemistry varied spatially and/or temporally. Nitrogen species were excluded from PCA analyses as their concentrations were below detection limit at several locations/timepoints and nitrate concentrations also strongly correlated with conductivity (Pearson’s R = 0.88, p<0.001) and phosphate concentrations (Pearson’s R = 0.47, p<0.001). No clear clustering of samples by location or timepoint was observed despite drinking water samples being obtained from variable source waters (Figure 1B). For instance, CAY, GUA, and MAN had higher mean and variable conductivities compared to other locations, which is consistent with source water type of these locations being groundwater. Phosphate, on the other hand, was relatively narrowly distributed across samples, and in lower concentration compared to other DW systems(Gooddy et al., 2015), with CAY location consistently higher than other PR locations.

**Figure 1:**
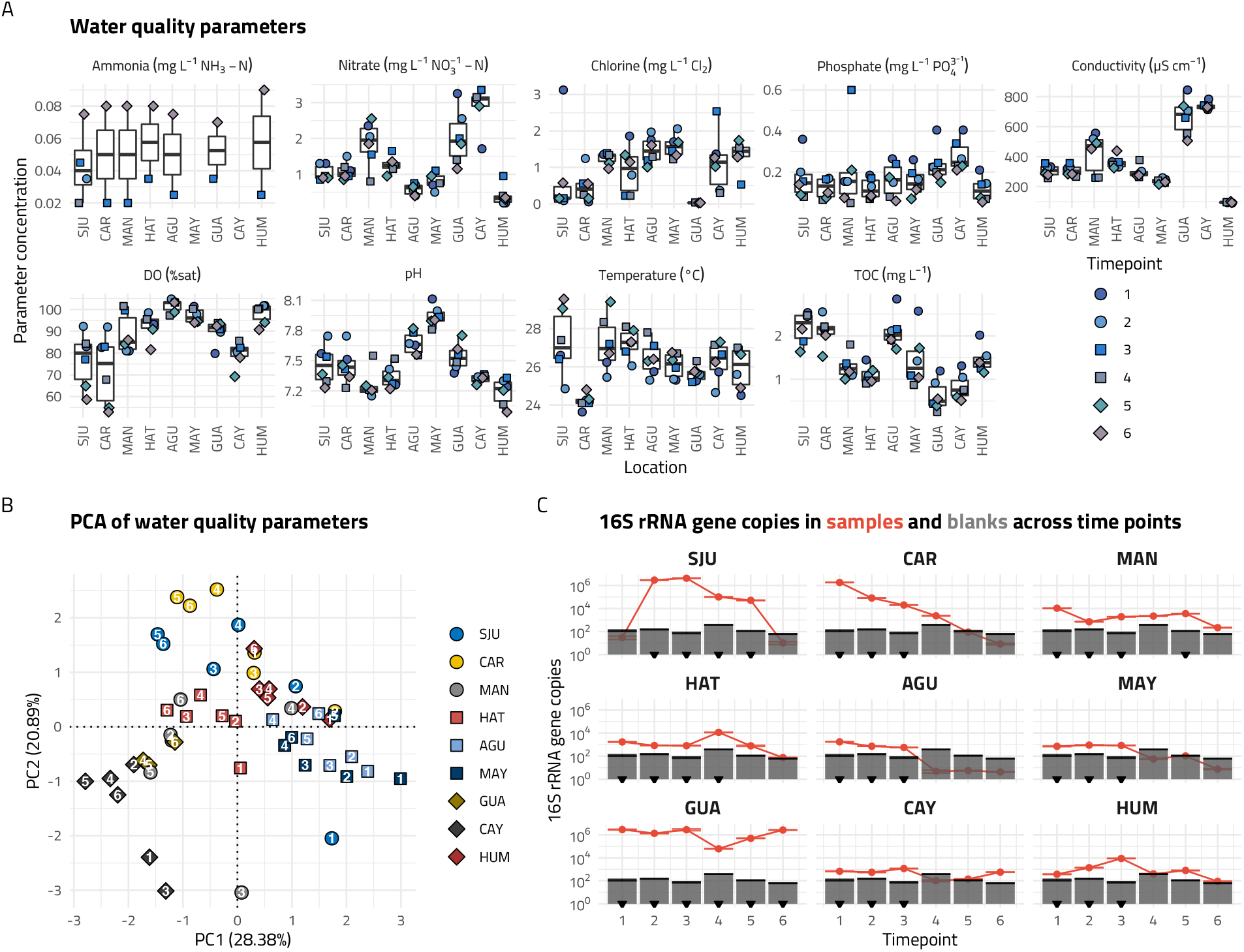
(A) Concentrations of water quality parameters for nine locations from December 2017 to October 2018. Shapes and colors correspond to timepoints. All nitrite measurements were below detection limit (BDL, not shown), for ammonia 36 measurements were BDL, and for chlorine 3 samples were BDL. (B) Principal component analyses of water quality parameters. Colors and shapes correspond to sampling location. Direct labels of timepoints within sampling location, in white. The PCA does not show a clear clustering between samples or timepoints. (C) Gene copies of 16S rRNA gene for samples and blanks. Red lines correspond to samples and grey bars correspond to blanks. Comparison between samples and blanks guided sample selection for sequencing. Inverted black triangles denote samples that were sequenced.

We quantified the abundance of 16S rRNA genes in all samples as a measure of bacterial concentrations (Figure 1C). The average PCR efficiencies for these qPCR assays was 90±2.9%. Volume normalized 16S rRNA gene copies (16S rRNA gene copies mL^-1^) ranged from 6.4×10^−2^-9.1×10^4^ copies mL^-1^. Independently regressing variables in the PCA as descriptors of log10(16S rRNA gene copies mL^-1^) for each location resulted in six significant (p<0.05) linear models. The goodness of fit for all models was relatively high with an average adjusted R^2^ of 0.704 ± 0.062, and significant associations between log10(16S rRNA mL^-1^) and DO in CAR, pH in HUM, temperature in CAR, MAY and GUA, and TOC in AGU. However, regressing parameters against log10(16S rRNA mL^-1^) for all locations only resulted in statistically significant associations with temperature (adj R^2^ = 0.067, p < 0.05) and total chlorine (adj R^2^ = 0.227, p < 0.001). A multiple linear regression model with all water quality parameters as descriptors of log10(16S rRNA mL^-1^) (n=51), indicated that total chlorine was the major driver associated with decreasing 16S rRNA gene concentrations (Adj R^2^ = 0.28, p<0.001, Figure 2A, Table S3). Chlorine concentrations measured in samples were comparable to those reported in other US studies(Stanish et al., 2016), except for GUA where it was either below detection limit (BDL) or very low for all timepoints. All samples, except timepoint 1 in SJU, had total chlorine concentrations below 3 mg L^-1^ (Figure 2B, Table S1). The negative relationship between bacterial load and chlorine concentration is particularly evident for SJU and CAR, where decreasing total chlorine concentration were associated with increased bacterial concentrations, relatively stable chlorine concentrations correspond to stable bacterial concentrations, and absence of chlorine shows high bacterial concentration, respectively (Figure 2B). Not surprisingly, these results suggest that maintaining chlorine residual is critically important for ensuring low bacterial concentrations which could be particularly challenging due to infrastructure damage from natural disasters. Some locations did exhibit significant variation in chlorine concentrations between December 2017 and February 2018 (e.g., SJU, CAR, HAT), with some of these variations associated with water main breaks (e.g., CAR in December 2017).

**Figure 2:**
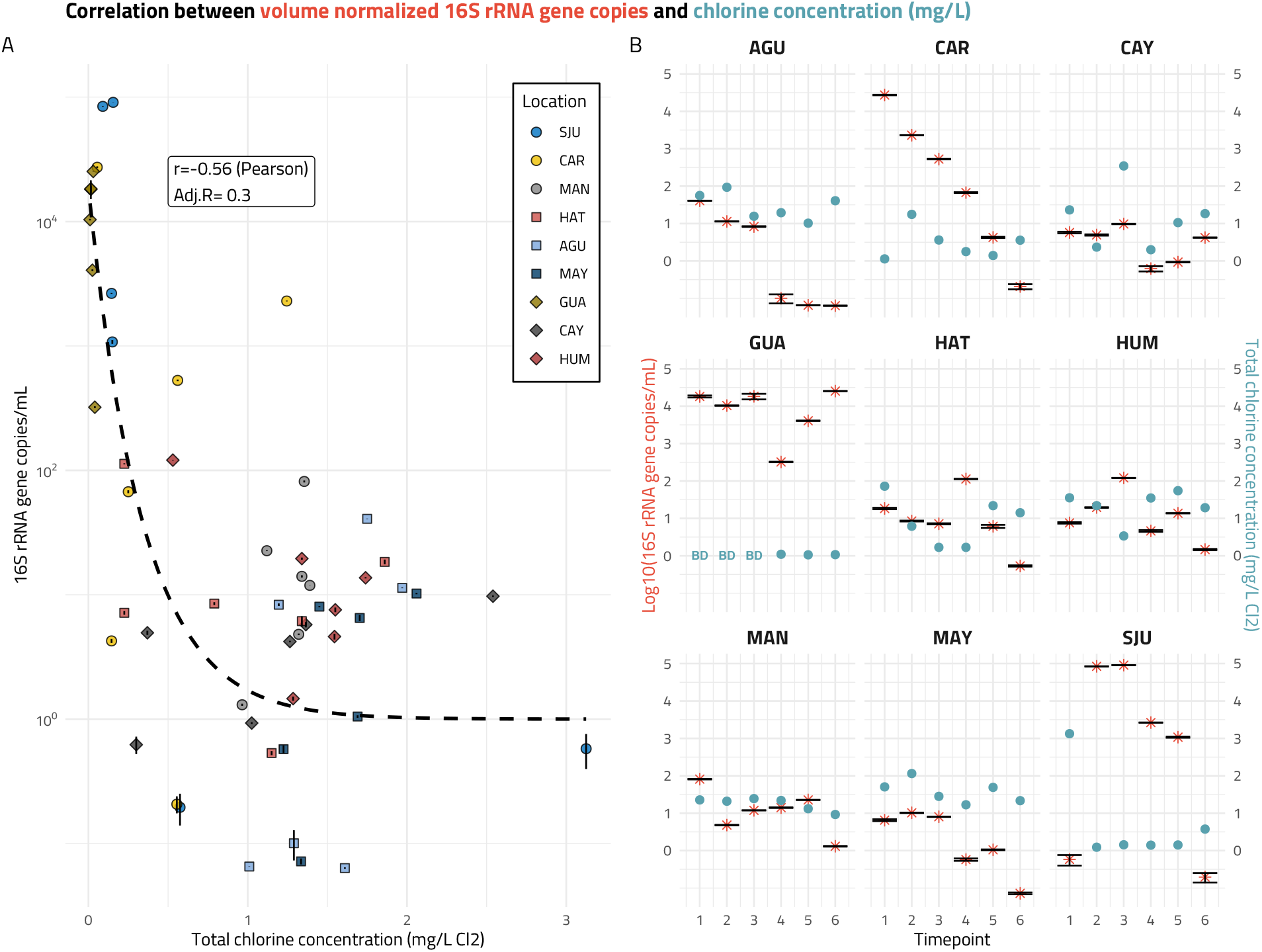
(A) 16S rRNA gene copies normalized by volume correlated with total chlorine concentration. Colors and shapes correspond to sampling location. Total chlorine concentration was the major driver among all recorded water quality parameters when all samples were aggregated in a multiple linear regression model. The dashed line corresponds to an exponential decay fit characteristic to inactivation of bacteria. (B) Double y-axis plot with the concentration of log 16S rRNA gene copies normalized by volume (left, red) and total chlorine concentration (right, blue) by time point faceted by sampling location. Chlorine concentrations were compliant with EPA regulation and bacterial concentrations were within range of typical drinking water systems. Total chlorine concentration was below detection (BD) at GUA first three timepoints. On a per location basis, clear examples of the negative effect of total chlorine can be observed (i.e., SJU, MAN, and GUA), however other parameters may drive bacterial concentrations in the remaining locations.

### 3.2 Microbial communities and metagenomes of PR samples were similar to those seen in other drinking water systems

Based on total bacteria qPCR results and comparison to blanks, a select number of samples per location were subjected to metagenomic sequencing (n= 33, Figure 1C, Table S4). The initial three sampling points for all locations were sequenced, unless their 16S rRNA gene copy numbers were below or equal to their matched controls. Further, any other sampling point 10-fold or higher 16S rRNA gene copy numbers than the highest observed in the controls for corresponding timepoint was sequenced as well. A total of 1.18 Gb raw reads were generated after quality filtering and 1.16 Gb reads were not mapped against UniVec, resulting in less than 2.6% of the reads being discarded for the majority of samples (n=31), only 2 samples (i.e. HUM_1, HUM_2) retained less than 92.5% of the raw reads (Table S4). Within sample diversity (Nd), as assessed by Nonpareil curves, ranged from 15.33-19.16 (Figure 3A, Table S5). These indices rely on redundancy of reads in a metagenome to estimate diversity of metagenome, with higher Nd corresponding to more diverse communities. The Nd values observed here are consistent with those seen in other chlorinated drinking water systems(Dai et al., 2020). The spread of observed Nd values within location was larger for GUA, HUM, and HAT, indicating higher temporal variation in diversity, while low variability in Nd values at CAR indicative of low temporal differences. Kruskal-Wallis test of Nd by location reveal significant differences in the median of at least one of the groups (p<0.05), however multiple hypothesis correction with Dunn test did not identify any significant pairwise differences. Significant and positive correlations were observed between Nd and pH at AGU (p<0.05), and Nd and DO at HUM (p<0.01), and significant negative correlations between HAT diversity and log10(16S rRNA mL^-1^) and CAY diversity and nitrate (p<0.05 for both locations).

**Figure 3:**
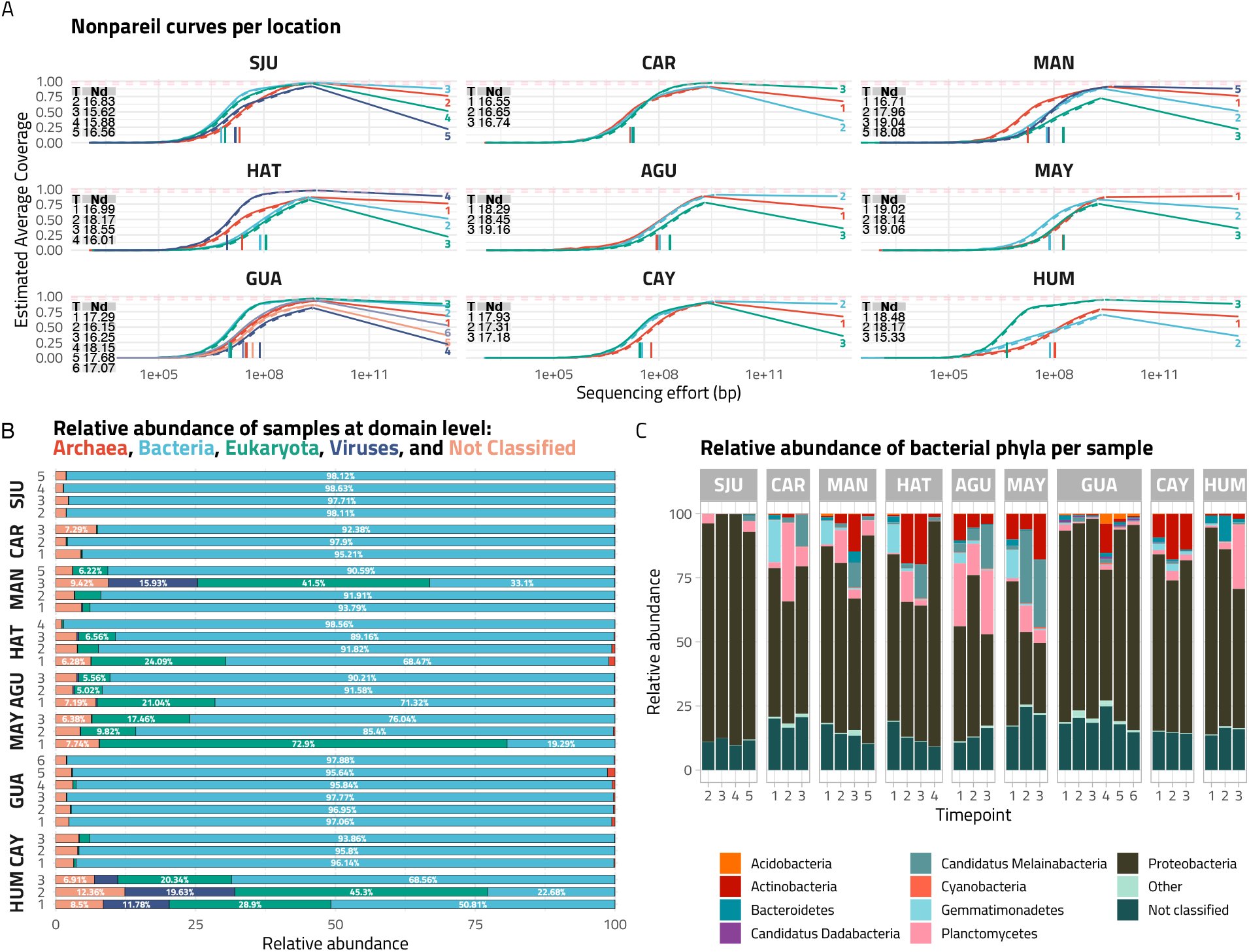
Diversity of microbial communities in metagenomes. (A) Nonpareil curves faceted by location. Each curve within facet denotes a sample directly labelled by timepoint. Respective Nonpareil diversity (Nd) is shown in table on the top left corner of each facet. The bars correspond to the sample estimated sequencing effort. Nd value indicates within sample community complexity in the sequence space. (B) Domain level relative abundances (RA) calculated with sample coverage information and CAT annotations. RAs greater than 5% are directly labelled. In some locations a large proportion of the samples were not classified, but Bacteria makes up the largest portion of most samples. (C) Relative abundances at Phylum level. Scaffolds classified to Bacterial domain were designated as 100%. Proteobacteria was the dominant phyla in the majority of samples.

The reads were subsequently assembled and scaffolds identified as potential contamination were removed as outlined in the materials and methods section. A summary of statistics for the 9 co-assemblies that were generated can be found in Table S6. CAT was used to annotate true scaffolds (i.e., scaffolds retained post contamination analysis) and coverage information allowed us to obtain per sample profiles (Figure 3B, Table S7). Bacteria generally constituted the largest portion in the samples from SJU, CAR, GUA, and CAY, with a mean relative abundance (RA) of 96.6±1.72%. In contrast, scaffolds of eukaryotic origin (18.5±19.5%) and unclassified scaffolds (5.62±2.85%) constituted a significant proportion of the community in MAN, HAT, AGU, MAY, and HUM. The RA of eukaryotic contigs is not consistent within locations, suggesting high temporal variation of the eukaryotic fraction in the systems. Despite highest eukaryotic contigs RA at timepoint 1 for AGU and MAY and timepoint 2 at HUM, water quality parameters at these sites had relatively low temporal variation. On the other hand, MAN had its highest RA at timepoint three when DO and phosphate concentrations were higher than mean values. In contrast with other sites with high contribution of eukaryotic contigs, MAN water source is groundwater. Phosphate concentration is associated with source water type and treatment processes (i.e., corrosion control). Douterelo et al.(Douterelo et al., 2018) compared metagenomic samples from sites with different source waters and saw dominance of bacteria and no significant differences in the RA of eukaryotes. Moreover Inkinen et al.(Inkinen et al., 2019) correlated the presence of phosphate concentrations with active eukaryotes in a chlorinated groundwater DWDS in contrast with surface waters. At the HAT site, the largest eukaryotic RA was seen at timepoint 1 consistent with highest chlorine concentration. Disinfection is a driver of microbial composition in DWS and it likely reduced the contribution of bacterial community in these samples, resulting in an observed increase in the RA of eukaryotes. We further investigate the eukaryotic component of the microbial communities identified through CAT with MetaEuk (Figure S1, Table S8 and S9). MetaEuk based classification indicated that free living amoeba (FLA) capable of supporting intracellular growth of opportunistic pathogens (e.g., *Vermamoeba, Acanthamoeba*, etc.) were transiently detected at low RAs at CAR, MAN, HAT, AGU, MAY, CAY, and HUM (Figure S2, Table S9). Waterborne parasites like Giardia and Cryptosporidium were not detected.

The coverage of scaffolds classified as bacteria was normalized by rpoB gene coverage in the respective samples to assess the bacterial community (Figure 3C, Table S10). The proportion of bacterial scaffolds not classified beyond the domain level ranged from 9.20-24.80% and patterns were consistent within location. Similar to previous studies, Proteobacteria was the dominant phylum in the majority of samples, ranging from 27.31-90.07%, with a mean of 64.2±16.6% for all samples. Actinobacteria and Planctomycetes were also detected in all samples. Actinobacteria is another group that is regularly detected in tap water(Hull et al., 2017). The Actinobacterial composition of MAN, HAT, AGU, MAY, and CAY tend to be higher relative to other locations, with an average RA of 9.10±6.95% in these samples, a mean 1.21±2.77% in other samples, and a global 5.27±6.61% RA. Planctomycetes is present at a RA greater than 1% in 72.72% of samples, but is predominant in CAR and AGU, with a mean RA of 13.41±15.08% and 20.58±7.24%, respectively, compared to a global average of 6.25±8.4%. On average, 81.2±9.61% of sample cumulative RA was not classified up to genus level. The dominant classified genera were Bradyrhizobium, Gemmata, Gemmatimonas, Hyphomicrobium, Methylobacterium, Mycobacterium, Novosphingobium, Pseudorhodoplanes, and Sphingomonas. We further compared the metagenomic assemblies recovered from the nine PR samples to other DWS (not impacted by natural disasters, i.e., undisturbed, Table S11), to assess if there were indications of significant deviation that could be attributed to HM. There was no clear clustering of metagenomes as shown by PCoA ordination of pairwise Mash distances including the nine co-assemblies from PR and 52 co-assemblies from other DWS (Figure 4A). Furthermore, there was no statistical difference between pairwise Mash distances grouped as PR vs other DWS and other DWS vs other DWS using permutational t tests (p>0.3, Figure 4B). This suggests that the differences in metagenomes between HM impacted and other DWSs are similar to those observed between other DWSs. Additionally, complete linkage clustering indicated that SJU, MAN, HAT, and AGU, and GUA and CAY clustered closely, both within and between each other; CAR did not cluster directly with another location and MAY and HUM were similar, but separate from the rest of the PR locations (Figure S3A). The respective Mash distances of early and late samples clustered identically as the previous analyses when leveraging coverage data and CAT classification of scaffolds to subset scaffolds pertinent to these categories (Figure S3B-C). This indicates that the metagenomes from the samples collected in PR were largely consistent with what would be expected from drinking water samples, irrespective of time of collection (i.e., December 2017 or October 2018).

**Figure 4:**
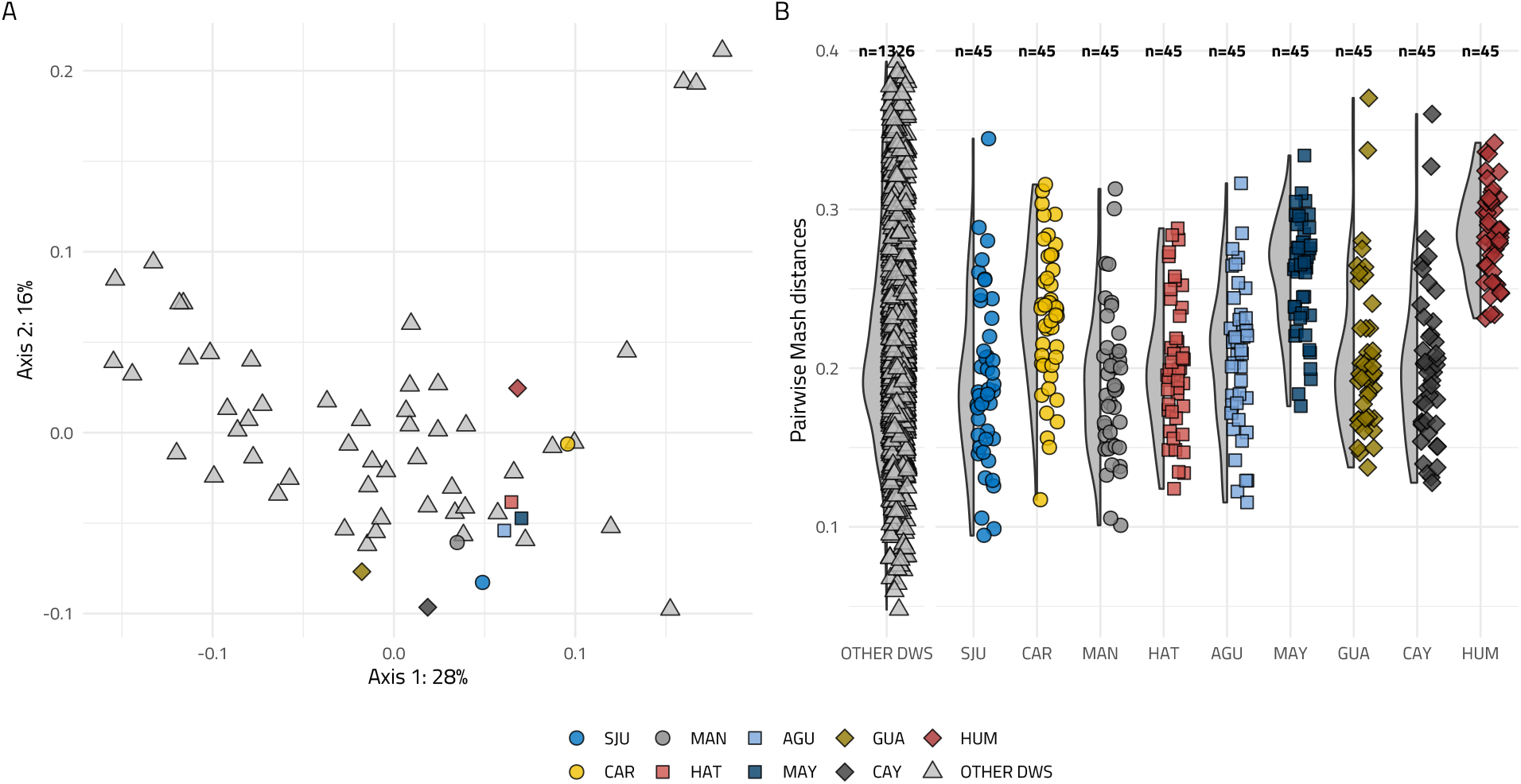
(A) PCoA ordination of Mash distances between DWS, including PR co-assemblies and reference DWSs. (B) Distribution of Mash distances prior to up sampling used for permutational t-test with group 1: other vs other and group 2: PR vs others. No significant differences were observed (p>0.3) between groupings. Colors and shapes correspond to PR locations or reference DWS.

### 3.3 Opportunistic premise plumbing pathogens were ubiquitous and detected at low concentrations

Genera that contain pathogenic species (i.e., Legionella, Leptospira, Mycobacterium, and Pseudomonas) were further investigated (Figure 5A) using CAT annotations with additional support from Kraken and/or Kaiju. While monitoring of indicator organisms and residual chlorine is part of the emergency response in the aftermath of hurricanes(Patterson and Adams, 2011), challenges with regulatory compliance were common in PR prior to HM and testing laboratories remained non-operational months after the hurricane. Further there was no systematic effort to monitor the prevalence of OPPPs (e.g., Legionella, Pseudomonas, and NTM). Heavy rain and flooding can severely impact water sources and as a result, distribution systems may increase the prevalence of pathogens in drinking water systems, leading to potential health risks.

**Figure 5:**
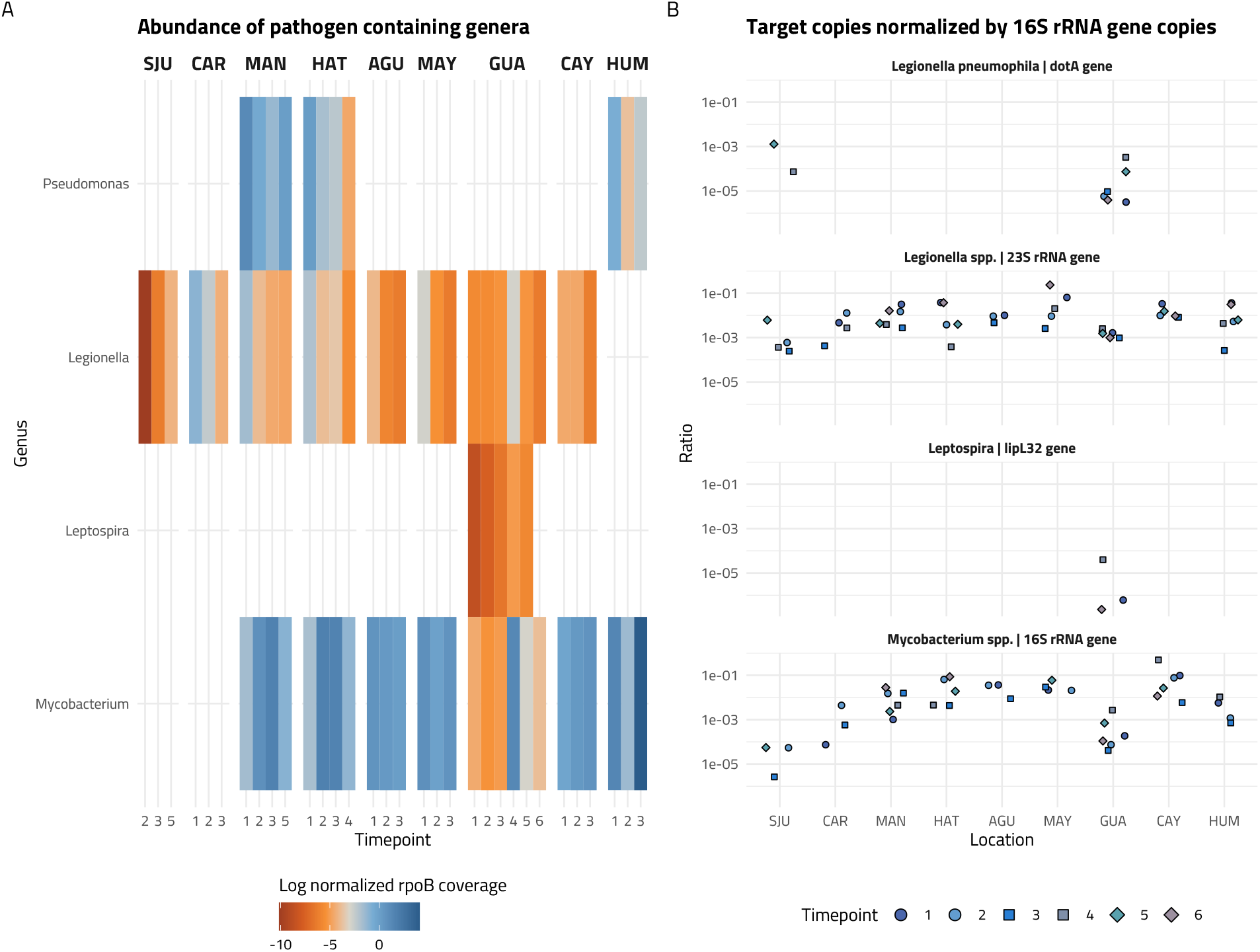
(A) rpoB normalized coverage of selected genera relevant to drinking water systems. Annotations are based on CAT classification and support from Kaiju or/and Kraken. (B) Ratio of copy numbers of targets (i.e. *Legionella pneumophila, Legionella spp*., pathogenic Leptospira, and *Mycobacterium spp*.) to 16S rRNA gene copies. Colors and shapes correspond to timepoints. Legionella spp. and Mycobacterium spp. are ubiquitous and more abundant than other targets. *Pseudomonas aeruginosa* or *Mycobacterium avium* were not detected in any samples.

Previous studies have reported the incidence of waterborne illnesses post hurricanes, including diseases with Legionella, NTM and Leptospira as causative agents(Maness, 2019; Shukla et al., 2018; Sutter and Sosa Pascual, 2018; Walker, 2018). Of the potential OPPP genera, Legionella and Mycobacterium were detected in most locations, while Pseudomonas was consistently detected in MAN, HAT, and HUM. Pathogenic Leptospira was only detected in GUA. There were statistically significant (Wilcoxon test, p<0.05) temporal differences in relative abundances of OPPPs for Legionella in MAN, Mycobacterium in HAT, MAY, GUA, and HUM, and Pseudomonas in HAT.

We used qPCR to quantify the abundance of *Legionella spp*. and *Mycobacterium spp*., while also conducting more targeted assays to detect and quantify the abundance of *Legionella pneumophila, Mycobacterium avium*, and *Pseudomonas aeruginosa*, and pathogenic species of the Leptospira genus (Figure 5B). The average qPCR efficiencies for these assays was ∼92.9%. Negative controls for each sampling campaign were included in every assay. *Mycobacterium avium* and *Pseudomonas aeruginosa* were not detected in any of the samples. The mean concentration for *Legionella pneumophila, Legionella spp*., pathogenic Leptospira, and *Mycobacterium spp*. in the samples was 0.3, 7.8, 0.01, and 1 copies mL^-1^, respectively. *Mycobacterium spp*. were observed in all locations with a general frequency of detection of 74%. *Mycobacterium spp*. were detected at every sampling timepoint in MAN, GUA, and CAY at very low concentrations (0.81±1.09 copies mL^-1^), while their concentrations were as high as 10 copies mL^-1^ at CAR and only detected in the first three timepoints. Consistent with metagenomic results, *Legionella spp*. was widely observed across all sampling locations at an 81% frequency of detection, with highest concentrations observed in SJU, CAR, and GUA. *Legionella spp*. concentrations decreased from 50 and 127 copies mL^-1^ at SJU and CAR, respectively to below 1 copy mL^-1^ from December 2017 to October 2018. *Legionella spp*. thrive in warmer temperatures(Lesnik et al., 2016), such as those in PR. Interestingly, concentrations *Legionella spp*. and *Mycobacterium spp*. are several orders of magnitude lower that what has been published in literature(Huang et al., 2021; Isaac and Sherchan, 2020; Ley et al., 2020; Liu et al., 2019) (i.e., 1-10^4^ copies mL^-1^). A potential reason for this could be over-chlorination in the systems, which had been reported in the aftermath of HM(Brown et al., 2018), including in early phase of sampling as this study showed.

The dotA gene assay to target *Legionella pneumophila* revealed low concentrations (i.e., 0.29±0.44 copies mL^-1^ for SJU and GUA locations. In SJU, *L. pneumophila* was detected in timepoints 4 and 5 only, while being detected at GUA at all timepoints. This is consistent with observations from other DWS where *Legionella pneumophila* was detected at low frequency and low concentrations(Lu et al., 2016; Wang et al., 2012).The LipL32 gene of pathogenic Leptospira were observed only in GUA and at timepoints 1, 4, and 6 (i.e., 0.01 copies mL^-1^, 5.6% frequency). The detection of Leptospira at this location was consistent with the detection of the genus Leptospira using metagenomics. Leptospira is not routinely reported in DWS, apart from the recent study by Keenum et al.(Keenum et al., 2021), possibly due to the efficacy of routine disinfection practices in the elimination of this pathogen(Wynwood et al., 2014). However, its importance has been highlighted in rivers and creeks when used as drinking water without proper treatment, particularly in situations of water scarcity, such as hurricanes(Keenum et al., 2021; Truitt et al., 2020). The presence of Leptospira in GUA is possibly exacerbated by the absence of residual chlorine at this location.

### 3.4 A small fraction of recovered metagenome assembled genomes were associated with pathogens

Metagenomes were co-assembled by location, binned, and manually refined with anvi’o. 105 bacterial MAGs were recovered after dereplication with dRep and quality filtering for completeness greater than 50% and percent redundancy lower than 10%. We identified one or more 16S rRNA in 39% of the MAGs. Further, we compared the differences in abundances between samples by accounting for MAGs read recruitment. Of these, 37% of the MAGs were detected in a quarter or more of the samples (Figure 6, Table S12). The PR MAGs were shared homology with 4.55% of a recently published JGI MAG collection based on ANI values ranging from 74.65-99.49%. JGI MAGs ecosystem categories represent aquatic (36.75%), human (31.31%), terrestrial (6.5%), built environment (5.03%), and wastewater (5%) environments. In contrast, the ecosystem distribution of the PR MAGs pairwise comparisons with JGI MAGs was comprised of aquatic (32.49%), terrestrial (20.57%), built environment (13.64%), plants (12.5%), and lab enrichment (4.95%) habitats. The aquatic ecosystem had the largest number of same species representation (20% of PR MAGs, Figure 6) with species boundaries level set at 83% cutoff threshold(Jain et al., 2018), and included 2 JGI MAGs also recovered from DWS.

**Figure 6:**
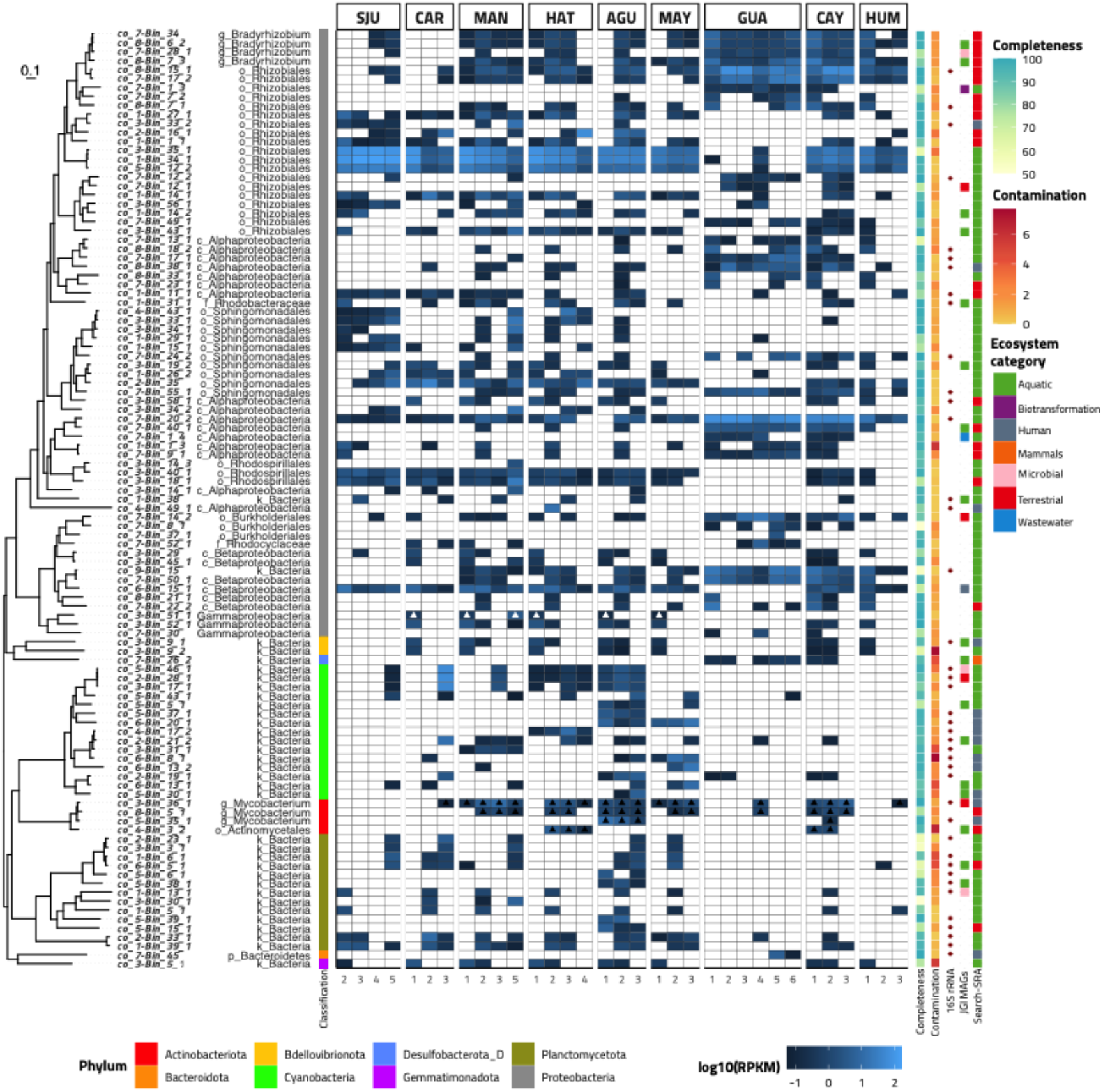
Metagenomic assembled genomes (MAG) information. From left to right: Phylogenomic tree of 105 recovered MAGs. Each branch is labelled by MAG code (italics) and marker lineage of MAG classified by GTDB-tk and annotated by color of phylum. Heatmap rows correspond to respective MAG and color gradient denotes log10(RPKM) values of MAGs detected in samples. A MAG was considered detected if ≥ 25% of its bases were covered by at least one read from the corresponding sample. Presence of color denotes detection of MAG and the blue gradient becomes lighter as abundance (i.e., RPKM) increases. Triangles in the heatmap highlight Pseudomonas spp. (white) and Mycobacterium spp. (black). MAG completeness ranges from 50-100% with color gradient from pale green to blue increasing value and contamination ranges from 0-10% with intense red corresponding to higher values. Red diamonds correspond to MAGs with one or more 16S rRNA gene detected. The highest ANI value between JGI MAGs and corresponding MAG from this study is colored according to ecosystem categories from JGI MAGs, if blank, no ANI above 83% was observed for that particular MAG. SearchSRA top environmental niche is depicted using the same color legend.

Despite this observation, the JGI MAG dataset suffers from lack of representation of MAGs assembled from DWS habitats (n=7). Therefore, a complimentary approach was used by mapping metagenomic reads from diverse ecosystems against the MAGs assembled in this study using the SearchSRA tool. This analysis indicated that the aquatic ecosystem was found to be the top environmental association for 63.8% of our recovered MAGs, the other top ecosystems were terrestrial (21.9%), human (13.3%), and mammal (1.0%) associated environments (Figure 6). However, if we consider the top four environments, aquatic ecosystem category is represented in all of our MAGs. There were statistically significant differences (ANOVA, p<0.001) between the proportion of reads mapping from each ecosystem category mapping to the PR MAGs, the only pairwise comparisons that were not statistically significant were terrestrial vs aquatic (Tukey’s, p=0.96) and mammals vs human environments (Tukey’s, p=0.06). Additionally, more than 13% of the aquatic ecosystem metagenomes were associated with DWS. Altogether, these analyses show that our MAGs are widely distributed in the environment, but are largely associated with aquatic and DWS associated environments.

The classification of resulting representative MAGs consisted of 64.8% Proteobacteria, followed by 14.3% Cyanobacteria, 12.4% Planctomycetota, and 3.81% Actinobacteriota. All of the Actinobacteria MAGs were classified as Mycobacterium. More than 50% of the MAGs were not classified to genus level. The most abundant genera among the MAGs included Hyphomicrobium (n=6), Bradyrhizobium (n=4), Gemmata (n=4), Mycobacterium (n=4), and Porphyrobacter (n=4). There was no relationship between environmental parameters and MAG abundance as assessed by Mantel statistic (r =0.133, p > 0.05) or constrained redundancy analyses. Three of the four Mycobacterium MAGs were classified up to species level and correspond to *Mycobacterium gordonae, Mycobacterium paragordonae*, and *Mycobacterium phocaicum*, all of which have been recovered from drinking water systems previously and are associated with infections in immunocompromised individuals(Shachor-Meyouhas et al., 2014). M. *gordonae* was more abundant (RPKM=2.27±2.77) and frequently detected (58%) than M. *paragordonae* (RPKM=0.77±0.41, 42% detection). Nevertheless, their abundance and frequency of detection was higher than for M. *phocaicum* (RPKM=1.8±1.9, 15% detection). Mycobacterium MAGs were not detected from SJU, and infrequently detected at CAR, GUA, and HUM. A single Pseudomonas MAG was recovered and classified as *Pseudomonas alcaligenes*. This MAG was only detected once and at the first timepoint at CAR, HAT, AGU, and MAY. At the MAN location, it was detected in the initial and final timepoint and at higher abundance at the final timepoint. *Pseudomonas alcaligenes* carries multiple antibiotic resistance genes, are considered opportunistic human pathogens and have been identified in previous literature characterizing drinking water systems, particularly in chlorinated systems(Jia et al., 2019; Ma et al., 2019).

## 4. Conclusions

This study characterized the microbial communities of nine locations in the aftermath of severe hurricanes (i.e., Irma and Maria) in a spatial-temporal yearlong survey using targeted and non-targeted molecular methods. Our results highlight that maintaining a disinfectant residual helps manage microbial concentration at the taps, yet sampling locations showed significant variation in the earlier timepoints. The estimated bacterial concentrations based on 16S rRNA gene abundance at the sampling locations were consistent with literature established values characterizing DWSs and generally decreased over time. Additionally, members of the microbial community were comparable to those found in other DWSs which were not impacted by natural disasters. Regardless of the ubiquity of some targeted OPPPs, such as *Legionella spp*. and *Mycobacterium spp*., they were present at low concentrations. Interestingly, pathogenic Leptospira was only detected at a single location and its presence could be associated with a lack of disinfectant residual at that site. A small fraction of metagenome assembled genomes were associated with potential pathogens, and other recovered MAGs represent previously reported taxa routinely found in drinking water systems. Altogether, the water disruptions (i.e., no water or intermittent supply) that were sustained after HM did not have a significant impact on the microbiological quality of drinking water in our study sites.

## Data availability

Metagenomic data is available on NCBI at Bioproject number: PRJNA718649 and the co-assemblies and metagenome assembled genomes are available on figshare at: https://doi.org/10.6084/m9.figshare.c.5414964.

## Acknowledgements

This study is supported by the United States National Science Foundation (NSF, CBET-1829754, CBET-1832756, IIS-1546428), the National Institute of Environmental Health Sciences (NIEHS) grants P42ES017198 and P50ES026049, and U.S. Environmental Protection Agency (EPA) grant R83615501. The authors acknowledge and extend their sincerest gratitude towards Lilliana Gonzalez, Jesus Lee-Borges, Perla Torres, Vibha Bansal, and Ezio Fasoli for their collaboration in sampling logistics.

