## Supplementary Figures for "Spatial-temporal targeted and non-targeted surveys to assess microbiological composition of drinking water in Puerto Rico following Hurricane Maria"

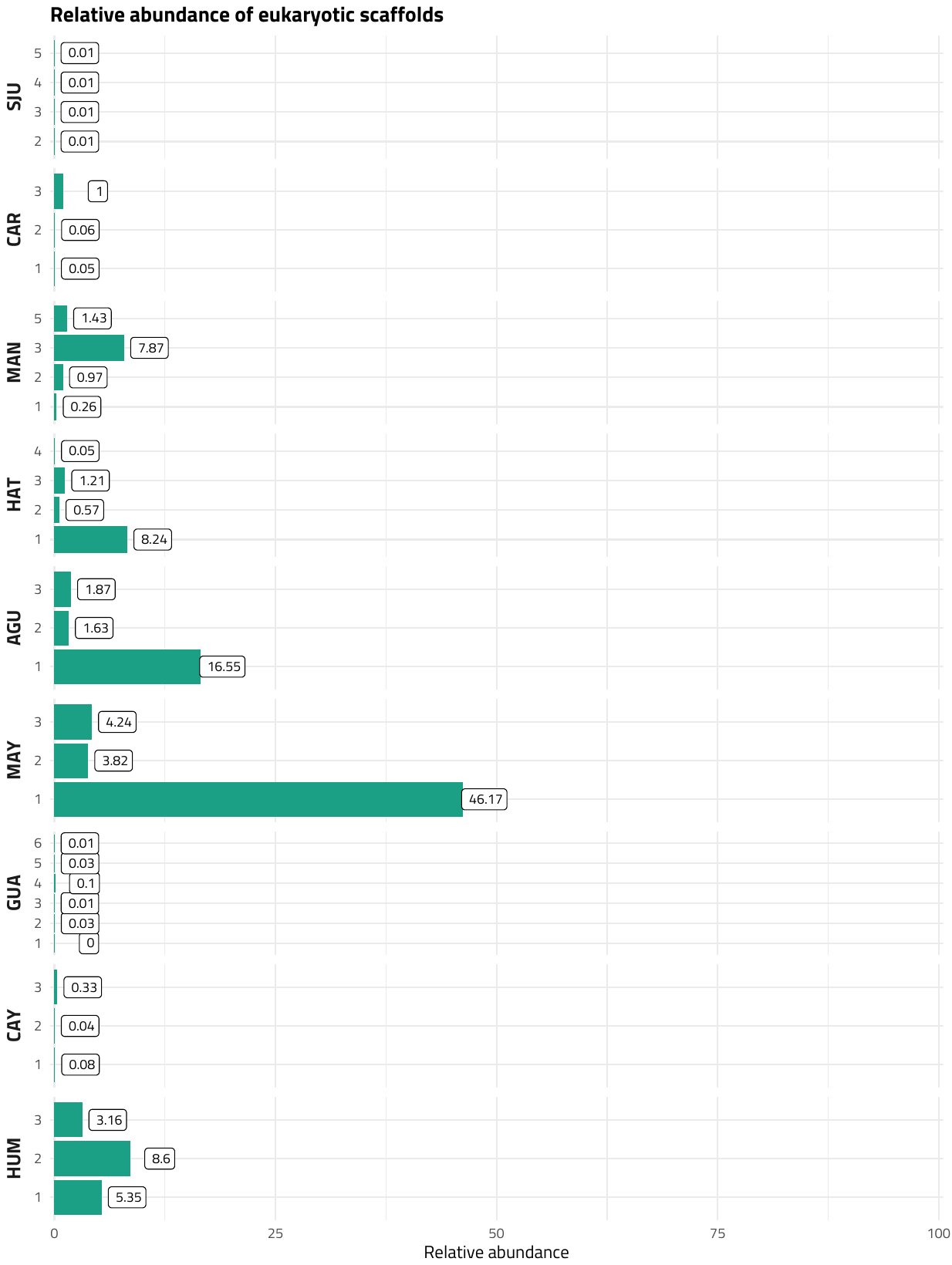

Figure S1: Relative abundance of eukaryotic contigs based on metaEuk classification.

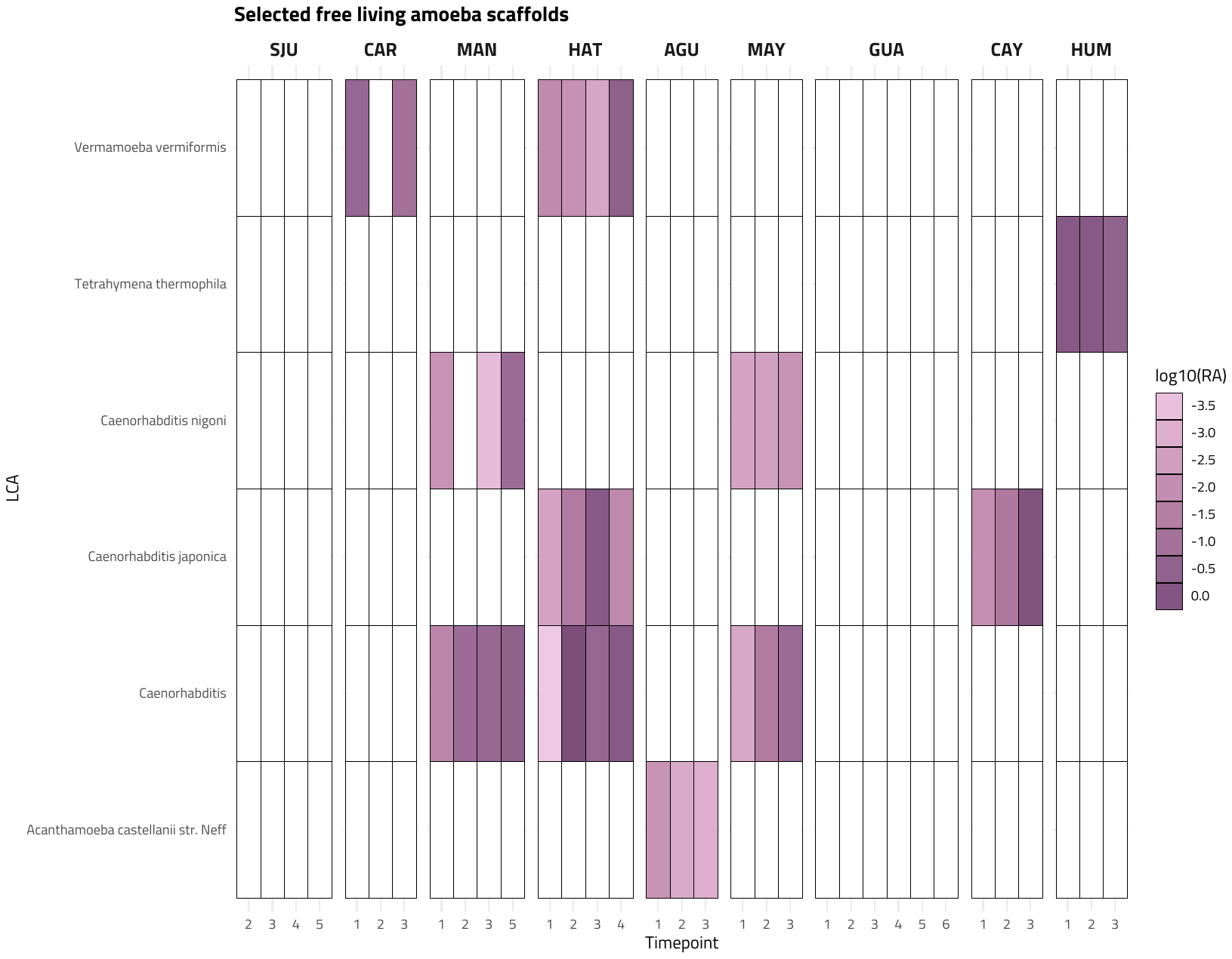

Figure S2: Relative abundance of selected free-living amoeba detected at the PR sampling locations based on MetaEuk based annotation of assembled scaffolds.

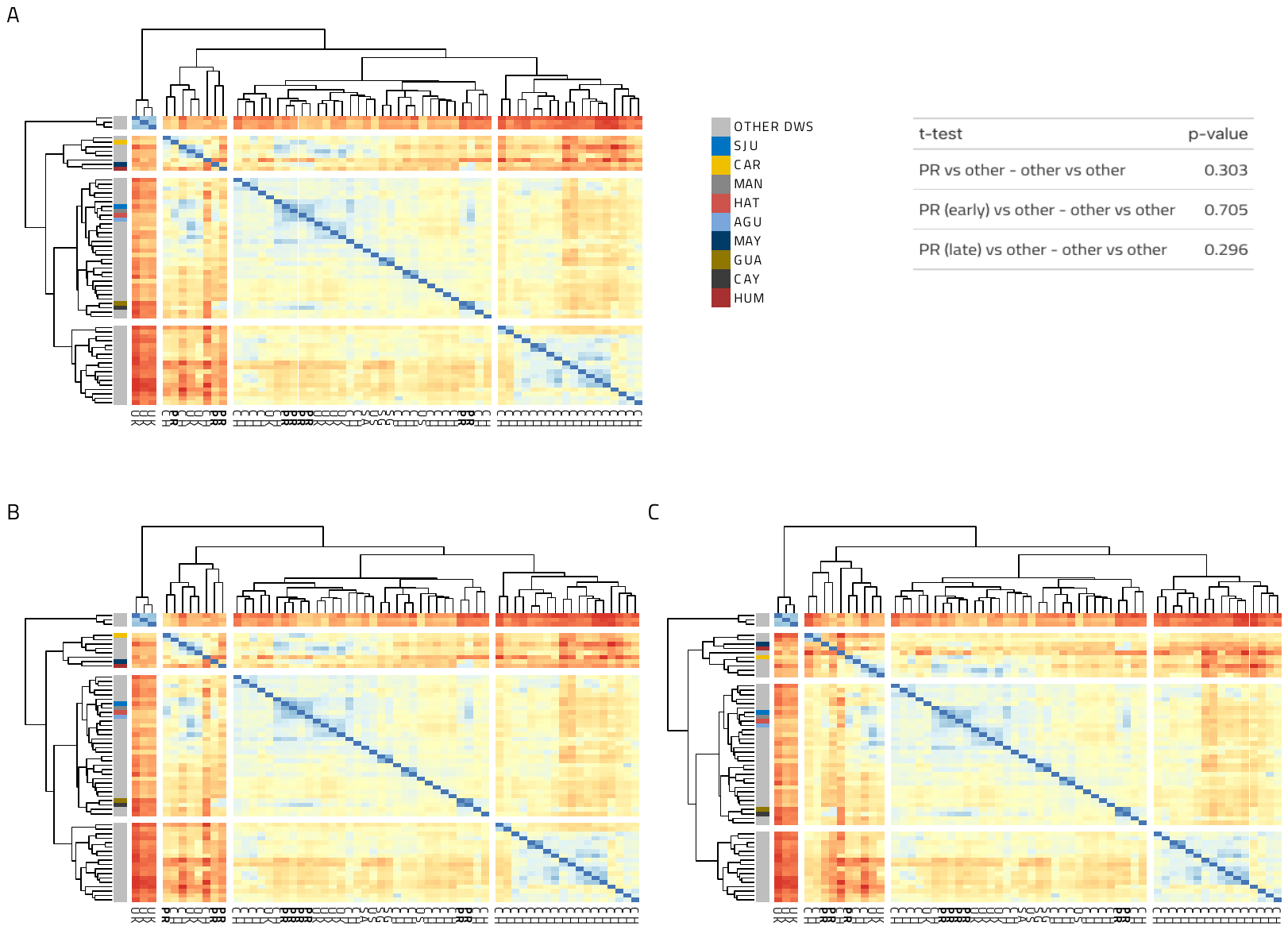

Figure S3: (A) Pairwise similarity of Mash distance between metagenomic assemblies pertaining from PR (this study) and other drinking water systems clustered by the complete linkage method using Euclidean distances. There is no clustering of metagenomes based on sampling country. MAY and HUM cluster separately from other locations. Furthermore, an identical clustering pattern is observed when sub-setting PR assemblies containing scaffolds from (B) early and (C) late timepoints, calculating similarity as outlined above.
